# Population Structure of Nation-wide Rice in Thailand

**DOI:** 10.1101/2021.05.09.443284

**Authors:** Phanchita Vejchasarn, Jeremy R. Shearman, Usawadee Chaiprom, Yotwarit Phansenee, Arissara Suthanthangjai, Jirapong Jairin, Varapong Chamarerk, Tatpong Tulyananda, Chainarong Amornbunchornvej

## Abstract

**Background:** Thailand is a country with large diversity in rice varieties due to its rich and diverse ecology. In this paper, 300 rice accessions from all across Thailand were sequenced to identify SNP variants allowing for the population structure to be explored.

**Results:** The result of inferred population structure from admixture and clustering analysis illustrated strong evidence of substructure in each geographical region. The results of phylogenetic tree, PCA analysis, and machine learning on population identifying SNPs also supported the inferred population structure.

**Conclusion:** The population structure inferred in this study contains five subpopulations that tend to group individuals based on location. So, each subpopulation has unique genetic patterns, agronomic traits, as well as different environmental conditions. This study can serve as a reference point of the nation-wide population structure for supporting breeders and researchers who are interested in Thai rice.

## Background

Rice (*Oryza sativa*) has been the main carbohydrate source in Thailand for more than 4,000 years [2], and Thailand has been a major rice exporter since 1851 [3]. Accelerated cultivar selection for specific environments is important for rice breeding programs. The long time period of rice domestication has yielded many rice cultivars with wide variation in physical traits, such as size, flowering time, grain quality, and yield, to name a few.

Thailand has large diversity in ecological systems [4]. In the north, most of the area is covered by mountains and tropical rain forests, while central Thailand consists of plains and fields that are prone to flood. In the north-eastern part, plateaus are the main type of area. In the south are tropical coastal regions and tropical islands. See Figure 1 for more details. According to Köppen climate classification [5], the south of Thailand is in the Tropical monsoon climate zone (Am), while the rest of the country is in the Tropical savanna climate zone (Aw/As).

**Figure 1.**
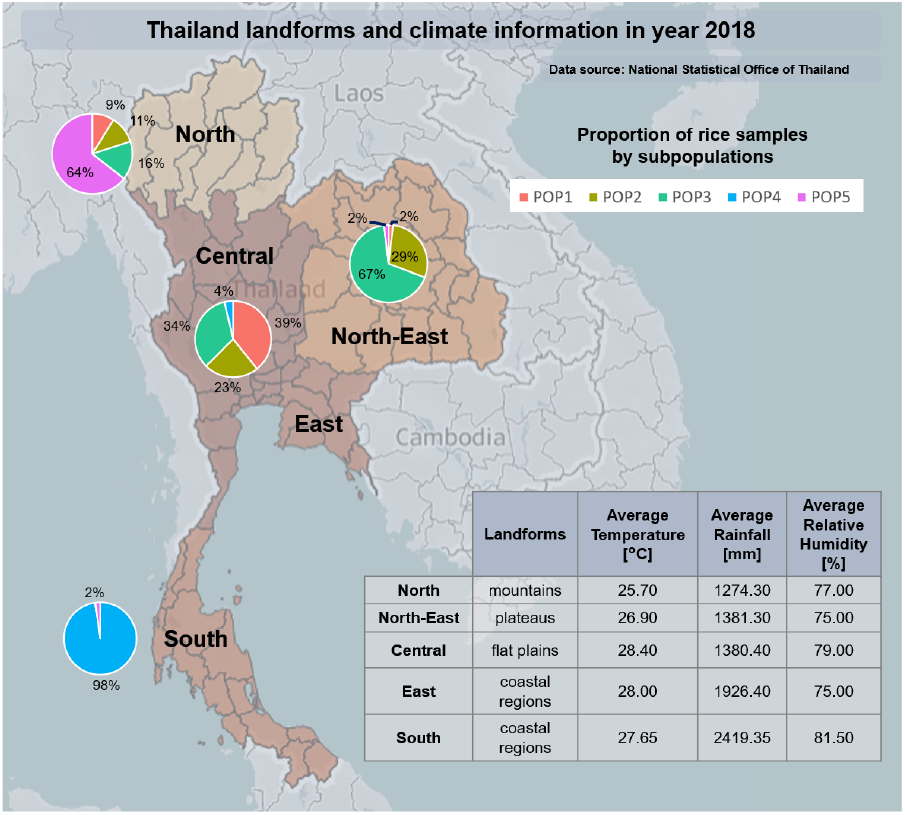
The environment of Thailand and the ratios of subpopulations in each area. The environment details are in the aspects of landforms, average temperature, amount of rain, and humidity in 2018 separated by regions (National statistical office of Thailand [1]). Each pie chart represents the ratio of each subpopulation members that have their known origin belong to the particular area. Note that there are no accessions for the east since it is not a rice cultivation area.

Due to the diverse ecology in Thailand, rice varieties need to be adapted to their intended growth region and there is some degree of association between genetic variation and geographical origin of Thai rice [6]. Moreover, there is a higher level of diversity in Thai rice accessions compared to selected rice accessions obtained from International Rice Research Institute (IRRI) germplasm based on InDel markers [4]. Limited data shows that Upland Thai rice forms a cluster of tropical japonica [7, 4, 8], while lowland rice forms indica clusters.

Understanding population structure and genetic diversity is an important step before Genome-wide association studies (GWAS) [9], which paves the way for studies of traits and functional gene investigation. Studies in population structure and genetic diversity of Thai rice have been conducted using different sets of rice varieties and molecular markers. Comparison of genetic diversity among 43 Thai rice and 57 IRRI rice varieties was investigated, using single-stranded conformation polymorphism (SSCP) InDel markers [4]. Additionally, 12 simple sequence repeat (SSR) markers were used to examine ongoing gene flow among three categories of rice variety in Thailand, including 42 wild rice varieties, 12 weedy rice varieties, and 37 cultivated rice varieties [10]. Recently, with a greater number of rice germplasm accessibility, 144 Thai and 23 exotic rice varieties were included to evaluate genetic diversity using SSR markers [7]. Another study assessed the population gene pool of 15 Thai elite rice cultivars using InDel markers [11]. It is worth noting that there are some limitations regarding access to a high number of accessions for each region of Thailand and the application of SNP markers to explore variation among Thai rice germplasms in these previous works.

To fill gaps in the literature, our study mainly focused on the population structure of 300 rice accessions, 277 of which are grown in diverse ecological systems in Thailand and 23 obtained from IRRI germplasm collection. We use SNP markers derived from the Genotyping-by-Sequencing (GBS) method to infer subpopulations. These accessions are a good representation of the nation-wide rice population structure.

## Results

### Population Structure

After clustering the 300 accessions, five subpopulations were found in the dataset. These five inferred populations generally group according to geological areas of rice accession cultivation.

Table 1 shows the origins of 300 accessions where the clusters of IRRI accessions were labeled according to the work in [12]. POP1 has a majority of indica accessions from Central Central Thailand. POP3 has a majority of indica accessions from Northeastern Thailand. POP2 represents rice accessions from both Northeastern and Central Thailand, suggesting it is an admixed population of the two. POP4 represents accessions from Southern Thailand. And lastly, POP5 represents japonica accessions from Northern Thailand. There are many accessions of indica from IRRI in POP1, which is consistent with POP1 being indica. The majority of japonica accessions from IRRI are in POP5, which includes the Thai japonica accessions. Additionally, a Chi-Square Test of Independence excludes the possibility that the origins and subpopulations in Table 1 are independent (36 dof, p-value ¡ 0.01). Hence, areas of origin and suppopulation in Table 1 are associated with each other.

**Table 1.** Origins of 300 rice accessions. There are 246 accessions from Thai known origins (north, north-east, central, or south), 31 accessions from Thai unknown origins, and 23 accessions from IRRI.

| Origin | Subpopulations |  |  |  |  | total |
| --- | --- | --- | --- | --- | --- | --- |
|  | POP1 | POP2 | POP3 | POP4 | POP5 |  |
| North | 4 | 5 | 7 | 0 | 29 | 45 |
| North-East | 1 | 14 | 37 | 0 | 1 | 53 |
| Central | 21 | 15 | 19 | 2 | 0 | 57 |
| South | 0 | 0 | 0 | 89 | 2 | 91 |
| IRRI indica | 7 | 1 | 0 | 0 | 0 | 8 |
| IRRI Tropical japonica | 1 | 2 | 0 | 0 | 2 | 5 |
| IRRI Aus | 0 | 3 | 0 | 0 | 0 | 3 |
| IRRI Temperate japonica | 0 | 0 | 0 | 0 | 4 | 4 |
| IRRI Aromatic | 0 | 0 | 0 | 0 | 3 | 3 |
| Unknown | 19 | 5 | 5 | 1 | 1 | 31 |
| Total | 53 | 45 | 68 | 92 | 42 | 300 |

A principal component analysis showed that PC1 separated the japonica population accessions (POP5) from the rest of the accessions, while PC2 separated the southern population accessions (POP4) from the central and northern accessions of indica (Figure 2). Lastly, PC3 separated the central indica accessions (POP1) from the northern indica accessions (POP3), with the accessions identified as admixed (POP2) joining the two, showing that the geographical separation is reflected in the genotypes of each accession. A phylogenetic tree was constructed and showed that the japonica population (POP5) was separated from the indica populations (Figure 2 (D)). Admixed accessions (POP2) were distributed among central (POP1) and northern (POP3) branches, supporting that POP2 is an admixed group of POP1 and POP3, while POP1, POP3, and POP4 were clearly separated from each other. Admixture analysis showed that POP1, POP3, POP4, and POP5 were grouped into different ancestors (different colors). POP2, however, had mixed ratios of ancestor A and B, which were the ancestors of POP1 and POP3. This confirms that POP2 is an admixed population of POP1 and POP3. POP1, POP3, POP4, and POP5 have high bootstrap support around 0.9, while POP2 has average support at 0.69 (Table 2). This is consistent with POP2 representing an admixed population of POP1 and POP3.

**Table 2.** Number of accessions and support of clustering assignment from bootstrapping for each population. The support number represents the likelihood that each cluster has the same set of members. Higher support implies a higher chance that cluster members are in the same population.

|  | Number of accessions | Average support |
| --- | --- | --- |
| POP1 | 54 | 0.98 |
| POP2 | 45 | 0.69 |
| POP3 | 67 | 0.92 |
| POP4 | 92 | 0.89 |
| POP5 | 42 | 0.99 |

**Figure 2.**
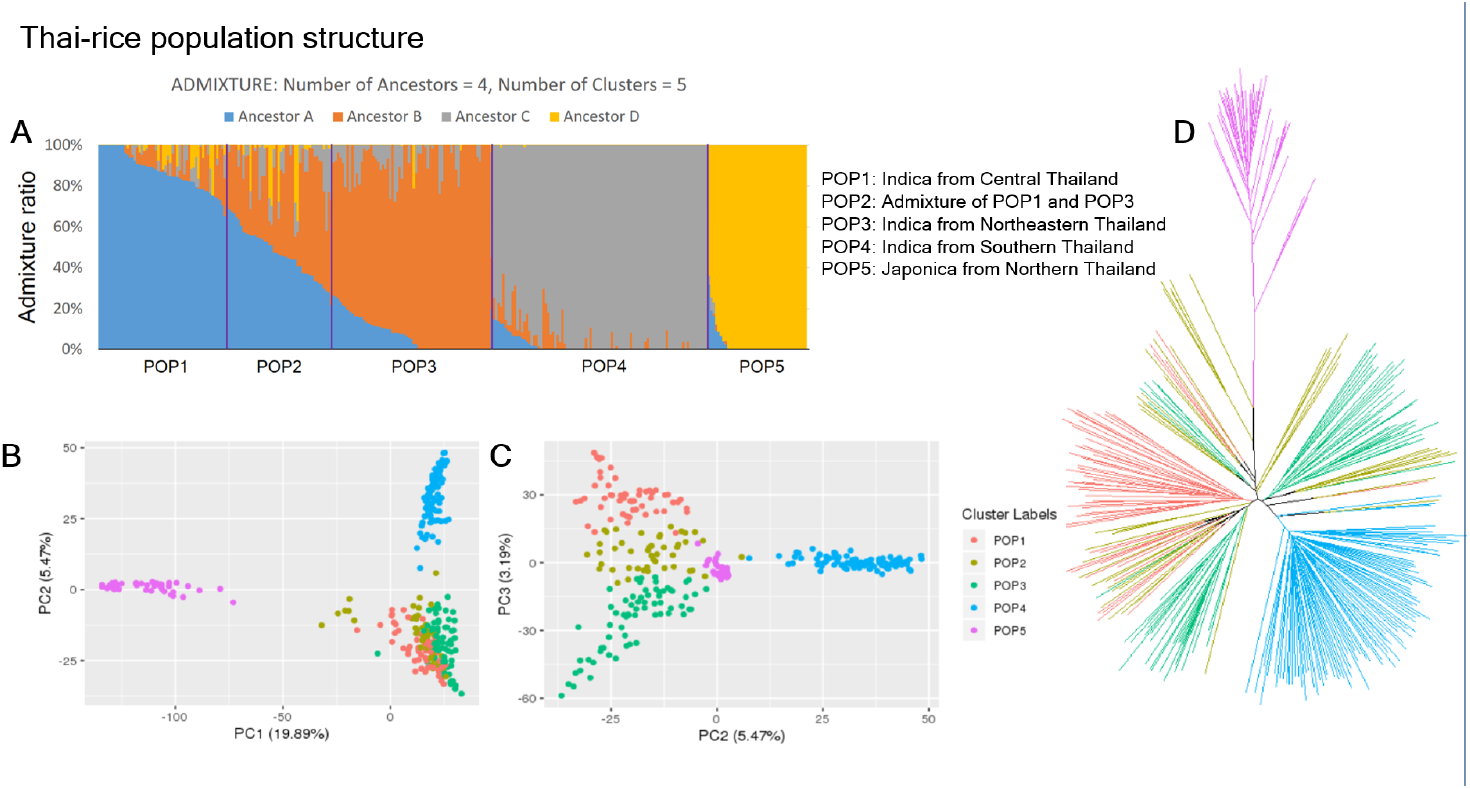
Population structure of 300 rice accessions inferred from 69,777 SNPs and 47,277 Indels. (**A**) Admixture plot of 300 rice accessions. The vertical axis represents an ancestry ratio of each accession. The horizontal axis represents individual accessions grouped by clustering analysis. Groups were assigned by clustering analysis on individual-admixture ratios. There are four ancestors (ancestor A - ancestor D) with five populations (POP1-POP5) inferred by clustering analysis. (**B**) The PCA scatter plot of first and second principal components (PCs) from a principal component analysis. (**C**) The PCA scatter plot of second and third PCs. Cluster colors were assigned according to ADMIXTURE clustering analysis results. The PC1 separates the japonica accessions (POP5) from the indica accessions. PC2 separates southern indica accessions (POP4) from central and northern accessions (POP1, POP2, and POP3). Lastly, PC3 separates central indica (POP1), from northern indica (POP3) with their admixture accessions appearing in between the two (POP2). (**D**) Phylogenetic tree of the 300 accessions, created by NJ tree, color coded according to the ADMIXTURE result.

The genetic distance of each population was estimated using *F*_*ST*_ between admixture ancestry populations, which is a widely-used measure of genetic variation among populations [13]. The *F*_*ST*_ (Table 3) shows that ancestor D, which was the ancestor of the japonica population (POP5), was the most distantly related.

**Table 3.** *F*_*ST*_ divergences between ancestry populations inferred by ADMIXTURE. *A* is an ancestor of indica (elite line), *B* is an ancestor of indica (modern variety), and *D* is the ancestor of japonica. By using a threshold of *F*_*ST*_ ≤ 0.3 to consider populations to have a similar type: either japonica or indica, *C* was assigned to be an ancestor of indica (landrace in southern part of Thailand).

| $F_{ST}$ | Ancestor *A* | Ancestor *B* | Ancestor *C* |
| --- | --- | --- | --- |
| Ancestor *B* | 0.178 | - | - |
| Ancestor *C* | 0.208 | 0.209 | - |
| Ancestor *D* | 0.480 | 0.497 | 0.507 |

The majority of accessions that formed POP4 were landraces from southern Thailand. These landraces were considered likely to be mostly indica, but there was no empirical evidence to support this. The *F*_*ST*_ values suggest that the ancestry of POP4 (C) was closer to ancestors A and B, which are indica, than to ancestor D, which is japonica. In addition, two indica accessions from the central region belong to the same cluster as the landraces. The members of POP4 cluster in PCA plots are the indica accessions rather than the japonica accessions (Figure 2). This shows that the landraces from southern Thailand are primarily of indica descent.

The 300 accessions were compared against 30 accessions of Thai rice selected from the 3,000 rice genomes project (3K RGP dataset) [14] that have areas of origin in Thailand using PCA (Figure 3). According to the result, indica accessions from the 3K RGP dataset are in POP1, POP2, POP3, and POP4, while japonica accessions from 3K RGP dataset are in POP5. These 3K accessions are consistent with the population groupings. An indica-japonica admixed variety from the 3K RGP dataset is placed between the area of japonica and indica in the PCA (Figure 3).

**Figure 3.**
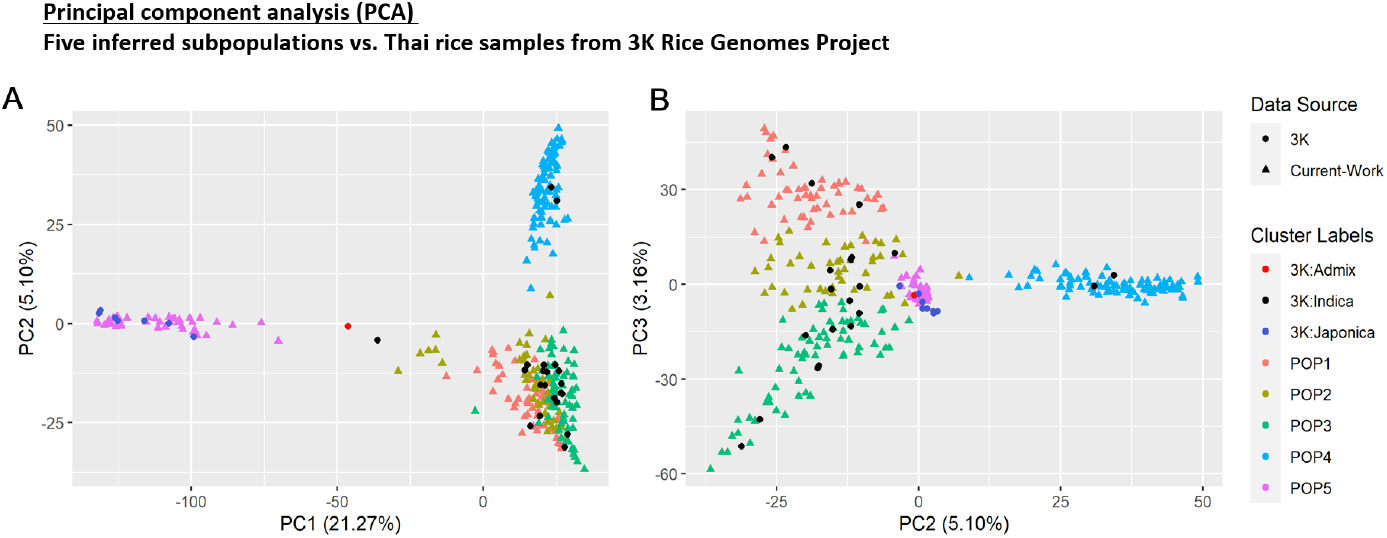
The Principal component analysis (PCA) for five inferred subpopulations and Thai rice accessions selected from 3K Rice Genomes Project (3K RGP dataset). (**A**) The PCA scatter plot of first and second principal components (PCs). (**B**) The PCA scatter plot of second and third PCs.

Additionally, many accessions from the Southeast Asian Indica (IND3) are grouped with POP4 (see Supplementary Table 7 for details regarding types of clusters in 3K RGP dataset).

### Agronomic traits of subpopulations

There are three agronomic traits that were measured for all accessions: days to flowering, grain length, and plant height.

The broad-sense heritability estimates 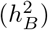 were 97.8% for plant height, 98.6% for grain length, and 93.58% for flowering time.

For the days to flowering trait, central indica accessions (POP1) flowered earlier than north-eastern indica accessions (POP3). The admixed population (POP2) had a flowering time roughly between that of POP1 and POP3, as expected. Southern indica accessions (POP4) have the latest flowering time of the 300 accessions investigated. Lastly, the japonica accessions (POP5) had a similar flowering time to POP1 (Figure 4 A,D).

**Figure 4.**
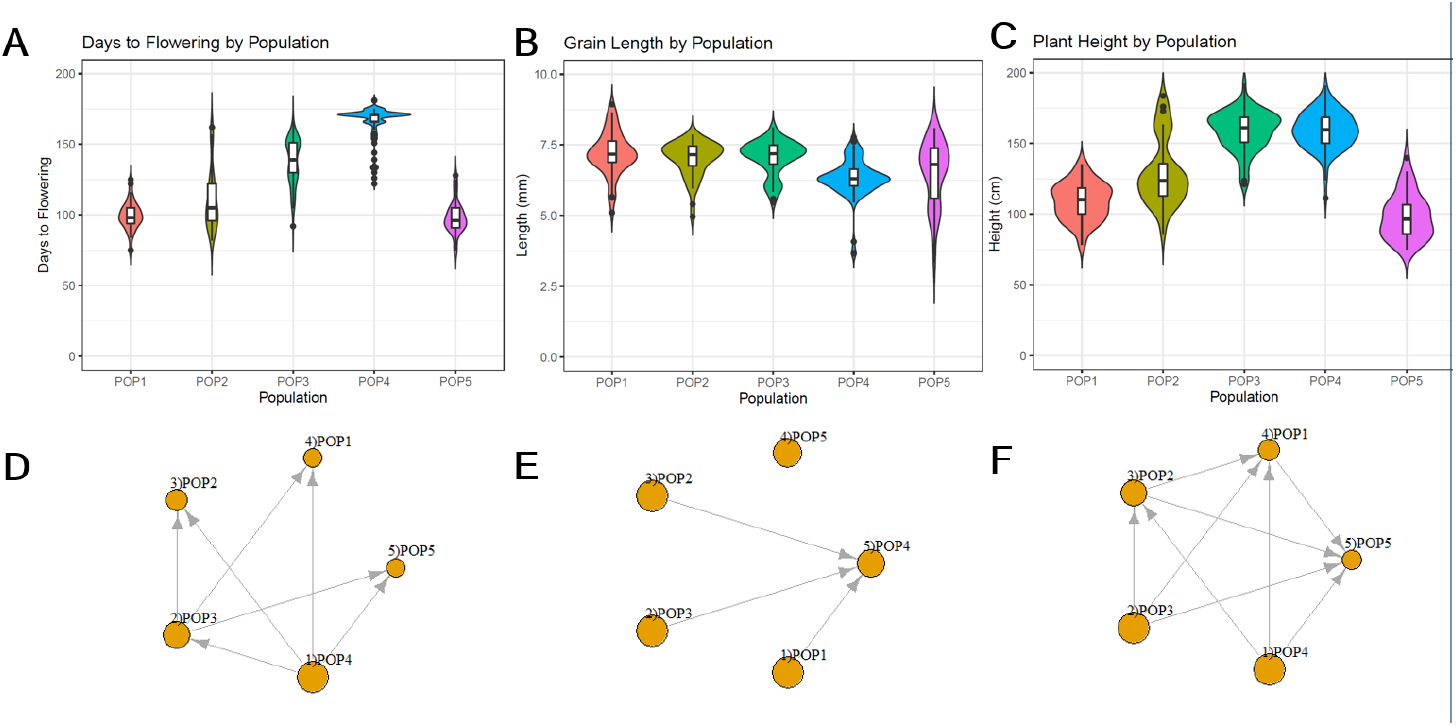
Subpopulation distributions of three phenotypes: days to flowering (A), grain length (B), and plant height (C). Domination graphs represent relationships between pairs of populations for days to flowering (D), grain length (E), and plant height (F). Arrow directions point from the population with a significantly higher phenotype value to the population with a lower phenotype value (with Mann Whitney test at *α* = 0.001).

**Figure 5.**
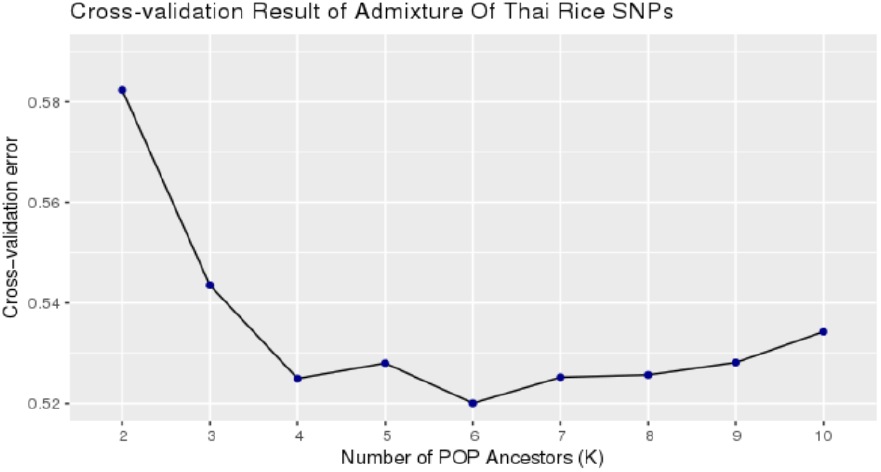
The Elbow method result that was used to find the optimal number of ancestors from ADMIXTURE. The optimal number is 4 ancestors.

**Figure 6.**
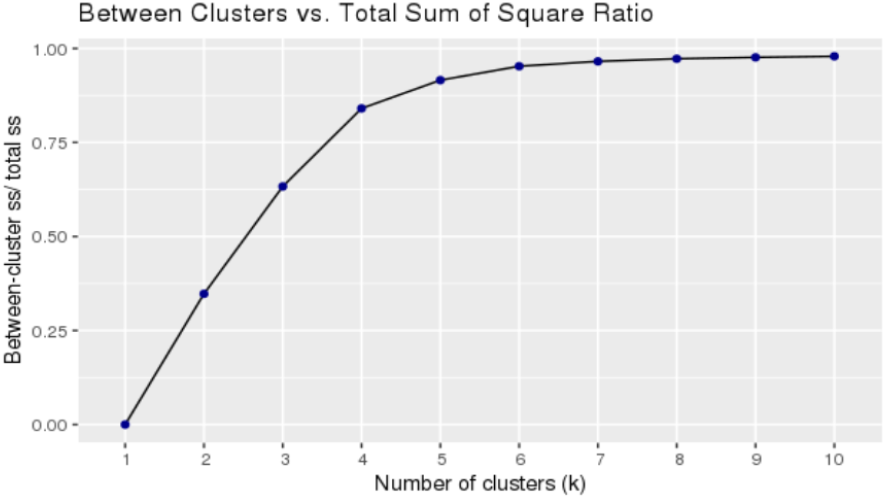
The Elbow method result that was used to find the optimal number of clusters for k-means clustering. The optimal number is 5 clusters.

The reason that rice in the central and northeast regions have different flowering times, even though the two areas have a similar latitude, is primarily because of distinct environment conditions in these regions to support multiple growing seasons per year. In central plain of Thailand, an irrigation system is well-managed and feasible for off-season rice cultivation, so farmers choose to grow short-duration rice varieties which can be harvested faster. Hence, there are more than one growing seasons per year in the central area. In contrast, the north-east has less rainfall and less access to water sources compared to the central region. Northeastern farmers tend to cultivate rice in one growing season per year and select for drought-tolerate varieties. This suggested different selection pressures for a days to flowering trait observed in this study.

For the grain-length trait, POP1, POP2, and POP3 have similar grain length, while POP4 has a significantly shorter grain length, and POP5 has high variation of grain length. This indicates that japonica (POP5) cannot be distinguished from indica (POP1 - POP4) by using the grain-length trait (Figure 4 B,E).

For the plant-height trait, ordering by ascending heights, the order is POP5, POP1, POP2, and POP3 / POP4. POP3 and POP4 have no significant difference in the height trait (Figure 4 C,F).

In the aspect of the association between phenotypes and known origins of accessions, with the Mann-Whitney test at *α* = 0.001, the results were as follows. The accessions from the south had significantly longer flowering time and significantly shorter grain lengths than the rest. The accessions from the north had significantly shorter flowering time than the rest. The accessions from the north-east had significantly longer plant height than the north. The accessions from the central area had significantly shorter plant height than the south. Hence, accessions can be separated roughly by these three traits, which implies that there are associations between traits and areas of origin of accessions. The potential cause of the difference in phenotypes might be the difference in landform and selection for crop use.

### Unique SNPs of subpopulations

A QTL analysis was used to identify SNPs with large variation in allele frequency between populations and 50-100 of the SNPs with the greatest allele frequency difference between populations were selected to train a random forest model to identify which population any given accession is from based on genotype. A total of 268 SNPs were selected (Supplementary Table 1).

Only POP5 had population specific SNPs that allowed for accurate population identification, this was not surprising as this population is japonica and the other populations are all indica (Table 4). The indica populations had too much allele sharing to allow for each accession to be accurately assigned to their population. The admixed population had the lowest rate of correct population assignment, while the other populations were all in the 80-90% range (Table 4.)

**Table 4.** The result of 10-fold cross validation based on 268 SNPs for population classification using Random Forest algorithm

|  | Precision | Recall | F1 |
| --- | --- | --- | --- |
| POP1 | 0.83 | 0.93 | 0.88 |
| POP2 | 0.76 | 0.62 | 0.68 |
| POP3 | 0.90 | 0.91 | 0.90 |
| POP4 | 0.97 | 0.98 | 0.97 |
| POP5 | 1.00 | 1.00 | 1.00 |

While a QTL analysis to identify population specific SNPs might be unconventional, it is well known that population stratification can result in false positives. In this particular case the populations in question are not discrete populations, but rather groupings of accessions that tend to correlate with location and have genetic mixing between groups.

The majority of SNPs most predictive for POP1 occurred on chromosome 1 in an interval between 21.6 and 22.5 Mb and an interval on chromosome 3 between 8.4 and 8.8 Mb. The majority of SNPs most predictive for POP2 occurred on chromosome 3 between 31 and 31.5 Mb with some small intervals on chromosomes 5, 6 and 7. There were 5 intervals of predictive SNPs for POP3 and several small intervals. Chromosome 3 had a interval from 27.59 to 27.65 Mb, chromosome 5 had an interval from 18.71 to 18.78 Mb, chromosome 6 had two intervals from 7.61 to 7.68 Mb and 11.02 to 11.06 Mb, chromosome 10 had an interval from 14.74 to 14.8 Mb. POP4 had the most distinctive allele frequencies with SNP intervals on chromosome 1 at 21.07 to 21.11 Mb, chromosome 2 at 5.32 to 5.35 Mb and 16.41 to 16.45 Mb, chromosome 5 at 23.71 to 23.84 Mb, and chromosome 11 at 2.7 to 2.8 Mb and 23.36 to 23.42. Of the 268 SNPs, there were 110 SNPs located in 75 genes, although the majority of these are predicted genes with no known function (Supplementary Table 2). There were 259 genes within the upstream and downstream intervals of the 268 predictive SNPs and most were predicted genes of unknown function (Supplementary Table 3).

## Discussion

According to the work in [4], upland Thai rice grouped into a japonica cluster, while rice from other regions formed an indica cluster, which is consistent with the population structure found in this work. Additionally, PCA analysis of rice accessions in this work compared against accessions from the 3K rice genome project confirmed that POP5 is japonica, while the rest of the subpopulations are indica.

All of the accessions of rice in this study possess unique traits that make them suited to their growing environment and type of farming. The types of environmental conditions range from the tropical monsoon climate in the south to tropical savanna in central Thailand and mountainous regions in northern Thailand. Grouping the accessions on genetic similarity tended to group accessions according to these environmental differences, which suggests that accessions in similar environments share the genetic variance that makes them suited to those environments.

The inferred subpopulation in the north is a japonica cluster (POP5). The other four inferred subpopulations are indica clusters in the central area (POP1), northeast (POP3), south (POP4), and the admixture of POP1 and POP3 (POP2). All inferred subpopulations were different and could be separated fairly well using 268 selected SNPs using Random forest classifier, with the exception of the admixed cluster (POP2). This implies that the inferred subpopulations were reasonably robust.

An interesting finding was that the most predictive SNPs for each subpopulation occurred within a few small intervals, rather than randomly spread throughout the genome, which may suggest a selection pressure, perhaps selecting for a trait that makes the accession better in the area it is grown. However, the subpopulation groupings are broad, each covering a quite diverse range of environments, and the allele frequencies between subpopulations have a large amount of overlap, so many of these regions could be due to chance rather than function.

Although the majority of genes within or nearby the SNP intervals have an unknown function, some interesting genes are functionally annotated, for example, *Os03g0262000, Os05g0203800, Os06g0677800*, and *Os09g0433650*. The gene *Os03g0262000*, is a homolog of *AtPIP5K1* that is induced by water stress and abscisic acid in *A. thaliana* [15]. *Os05g0203800 (OSMADS58)* is identified as a rice C-class MADS box gene which plays a crucial role for flower development [16], [17], [18], [19], [20]. *Os06g0677800 (OsARF17)* encodes a rice auxin response factor (ARF) involved in plant defense against several different types of plant virus [21], and functions in leaf inclination regulation [22] and tiller angle modulation [23]. *Os09g0433650* is located on chromosome 9 and associated with rice grain shape [24]. The roles of these candidate genes identify a potential relationship between predictive SNP markers and differences in agronomic traits found in the inferred subpopulations which could be further investigated.

## Conclusion

Thailand is a country with large diversity in rice varieties due to its rich and diverse environment. In this paper, 300 rice accessions (277 rice accessions from all across Thailand and 23 IRRI rice accessions) were sequenced to identify SNP variants allowing for the population-structure to be explored. The inferred population structure from admixture and clustering analysis illustrated strong evidence of substructure for each geographical region. The results of phylogenetic tree, PCA analysis, and machine learning on SNPs selected by QTL analysis also supported the inferred population structure. Moreover, by using only 268 SNPs, a random forest classifier was able to classify individuals for four out of the five subpopulations with reasonably high accuracy, the admixture population was the exception. This shows that these subpopulations are unique enough to be distinguished by a small number of SNPs. A unique ecological system where rice is grown might play a key role in this uniqueness. The 268 SNPs may be used as markers of these subpopulations for future studies. This study can serve as a reference point of the nation-wide population structure for supporting breeders and researchers who are interested in Thai rice. Finally, the dataset of 300 rice accessions is available at [25].

## Methods

### Plant material

The panel used in this study is composed of 300 Thai rice accessions representing diversity in phenotype, agro-ecosystem, and geographic origin: northern, northeastern, southern, and central region of Thailand. Detailed information regarding the accessions is reported in Supplementary Table 4.

### Plant cultivation

The study was carried out in the wet season of 2018 at Ubon Ratchathani Rice Research Center (URRC) of Ubonratchatani province, Thailand (15°19’55.2”N, 104°41’27.9”E). Seeds of the 300 rice accessions were germinated in a wet seedling bed on 16th June 2018. The seedlings were transplanted in a puddled field at 30 days after sowing (DAS) in 80 × 380 cm plots (5 rows x 20 plants). Fertilizers were applied as follows: 50 kg/ha N, 50 kg/ha P_2_O_5_, 25 kg/ha K_2_O at 10 days after transplanting; and top-dress with 10 kg/ha N at 30 days after transplanting. The experimental field was managed according to normal agricultural practices regarding crop protection and paddy water management. The mean air temperature ranged from 24.5 to 31.7°C. The highest and lowest relative humidity recorded during the experiment was 93.6 to 65.7%. No extremely high temperature or extremely low relative humidity was recorded, therefore heat stress was not a cause that affected growing and/or fertility conditions. Flowering time (days to flowering after sowing, DTF) was recorded when 50% of the plants in each plot had flowered. At maturity, the five plants in the middle position of each plot were selected for assessment of plant height and grain length. The broad-sense heritability 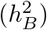 was calculated as 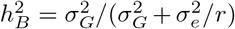, where 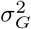 represents genetic variance, 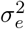 represents residual variance and *r* is the number of replicates.

### Genotyping by sequencing and variance calling

The genotypic sequences were generated from Ion S5™ XL Sequencer (Thermo Fisher Scientific). The data were obtained as BAM files. The ApeKI enzyme was used for genomic DNA digestion to prepare the DNA libraries for each accession. E-Gel™ SizeSelect™ agarose gels (Invitrogen) were used to select DNA fragments for 250–300 bp. Fastq files were created from BAM files using Samtools v1.9 [26]. Then, reads were mapped to the japonica reference genome using Burrow–wheeler aligner (BWA) v0.7.17 [27] and SAMtools. Variants were called using using GATK v4.1.4.1 [28].

### Population structure analysis

#### Numerical genotype function

Genotype was converted into a numerical value, such that homozygous reference allele was 1.0, homozygous alternate allele was 0.0, and heterozygous was 0.5 using TASSEL [29]. The SNPs were filtered to have a minimum allele frequency of 0.05 and a minimum call rate of 70% per SNP. The SNP number reduced from 3,366,491 to 117,054 sites after filtering.

#### Admixture analysis

Numerical genotypes were used to create .ped, .map and .bed files for ADMIXTURE [30] analysis to estimate ancestry ratios of all individual accessions. The optimal number of ancestors was found to be four by the Elbow method. The *F*_*ST*_ values where also calculated by ADMIXTURE [30].

#### Clustering analysis

The ancestry-ratio vectors of each SNP were used for data clustering. The individual assignments of clustering were inferred by applying a k-means clustering approach [31] in the R software package [32]. The Elbow method was applied to infer the optimal number of clusters based on Between-cluster and Total Sum-of-Square (BCTSS) Ratio. The BCTSS ratio represents a ratio of difference of distance from individuals to their cluster centroid between current clustering assignment compared to single cluster assignment. The optimal number *k*\* of clustering assignment should reduce BCTSS ratio significantly compared against *k*\* − 1 and *k*\* + 1 cases.

A 10,000 iteration bootstrap approach [33] was deployed to estimate the support of clustering assignment of each population. The clustering assignment that maximized BCTSS ratio with the optimal k along with the support of assignment from bootstrap were used to represent the subgroups of the population.

#### Principal components analysis

PCs were generated from numeric genotype data using TASSEL [29].

#### Phylogenetic tree construction

A phylogenetic tree was generated by Neighbor-Joining method [34] using the numerical genotype data in TASSEL [29].

#### Domination graphs inference

Domination graphs, which represent relationships between pairs of populations for three phenotypes, were inferred using EDOIF package [35]. For each phenotype, nodes of the domination graph are subpopulations while there is an edge from a population with a significantly higher phenotype value to a population with a lower phenotype value. The Mann Whitney test was deployed to infer edges of a domination graph with *α* = 0.001.

#### Population specific SNPs

We investigated the potential of identifying SNPs that were specific to each population identified by the admixture analysis. These groupings can include a large number of accessions and the accessions have varying levels of relatedness, which means varying levels of SNP sharing occur within and between populations, so a large number of SNPs would be required to discriminate between populations. The variants were filtered to select for bi-allelic SNPs where all accessions were homozygous and a series of Quantitative trait locus (QTL) analyses were performed to identify the most discriminatory SNPs. The phenotype for each QTL analysis was set as a binary trait of ‘same population’ or ‘other populations’ using the population groupings identified by the admixture analysis. A separate QTL analysis was performed for each population and the SNPs with the highest LOD score and largest allele frequency difference were taken as being the most predictive for that population. These SNPs were then used to train a random forest model [36] using the R randomForest package [37] and the R caret package [38]. Gene information from the GFF was overlaid on the SNP data to identify any population discriminatory SNP that was within a gene. In addition, genes within intervals of closely spaced predictive SNPs were also investigated.

#### Population classification

We deployed machine learning data classification to investigate whether the set of population specific SNPs we selected can be used to discriminate between the five populations. We used 10-fold cross validation [39], which is a technique in machine learning to measure the performance of prediction from a set of classifiers. We used a random forest model [36] as the main classifier in the analysis training on the 268 selected SNPs to classify the five populations of 300 rice accessions. A true positive (TP) is when the predicted population was the same as the ADMIXTURE derived population. The false positive (FP) count is the incorrect inclusion of an accession into a subpopulation and the false negative (FN) count is the incorrect exclusion of an accession out of a subpopulation, calculated per subpopulation. The precision is the ratio of the number of TP cases to the sum of TP and FP cases. The recall is the ratio of the number of TP cases to the sum of TP and FN cases. The F1 score is calculated from precision and recall as follows.

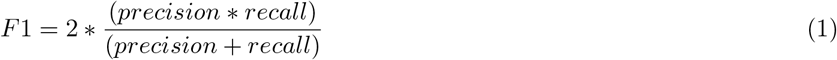

## Abbreviations

BCTSS: Between-cluster and Total Sum-of-Square ratio;
GWAS: Genome-wide association studies;
GBS: Genotyping-by-Sequencing;
F1: F1 score or F measure;
*F*_*ST*_: Genetic differentiation;
IRRI: International Rice Research Institute;
MAF: Minor allele frequency;
PCA: Principal component analysis;
POP1: Indica subpopulation originated from central part of Thailand;
POP2: Admixed subpopulation of north-eastern and central indica subpopulations;
POP3: Indica subpopulation originated from north-eastern part of Thailand;
POP4: Indica subpopulation originated from southern part of Thailand;
POP5: Japonica subpopulation originated from northern part of Thailand;
QTL: Quantitative trait locus;
SNP: Single nucleotide polymorphism;
SSCP: Single-stranded conformation polymorphism;
SSR: Simple sequence repeat.

## Declarations

### Ethical Approval and Consent to participate

Not applicable

### Consent for publication

Not applicable

### Availability of supporting data

Yes, please contact the corresponding author for the request of data access.

### Author’s contributions

P. Vejchasarn: Conceived and designed the experiments; Performed the experiments; Contributed reagents, materials, analysis tools or data; Wrote the paper. J.R. Shearman: Contributed reagents, materials, analysis tools or data; Analyzed and interpreted the data; Wrote the paper. U. Chaiprom: Contributed reagents, materials, analysis tools or data; Analyzed and interpreted the data; Wrote the paper. Y. Phansenee: Performed the experiments; Analyzed and interpreted the data; A. Suthanthangjai: Performed the experiments; Analyzed and interpreted the data; J. Jairin: Conceived and designed the experiments; Analyzed and interpreted the data; V. Chamarerk: Analyzed and interpreted the data; Wrote the paper; T. Tulyananda: Analyzed and interpreted the data; Wrote the paper; C. Amornbunchornvej: Contributed reagents, materials, analysis tools or data; Analyzed and interpreted the data; Wrote the paper.

## Acknowledgements

We also would like to thank Dr. Varapong Chamarerk, Mr. Ekkarat Kaewnango Mr. Apichart Noenplab, and Mrs. Kanthita Suangtho for supporting this work in many aspects.

## Supplementary

## References

1. (NSO), T.N.S.O.: Thailand Environment Statistics 2020. International series of monographs on physics. Thailand’s National Statistical Office (NSO), Bangkok (2020). http://service.nso.go.th/nso/nsopublish/pubs/e-book/Thailand_Environment_2020/files/assets/common/downloads/publication.pdf

2. Weber, S., Lehman, H., Barela, T., Hawks, S., Harriman, D.: Rice or millets: early farming strategies in prehistoric central thailand. Archaeological and Anthropological Sciences 2(2), 79–88 (2010). doi:10.1007/s12520-010-0030-3

3. Siamwalla, A.: A history of rice policies in thailand. Food Research Institute Studies 14(1387-2016-115909), 233–249 (1975)

4. Chakhonkaen, S., Pitnjam, K., Saisuk, W., Ukoskit, K., Muangprom, A.: Genetic structure of thai rice and rice accessions obtained from the international rice research institute. Rice 5(1), 19 (2012)

5. Köppen, W.: Die wärmezonen der erde, nach der dauer der heissen, gemässigten und kalten zeit und nach der wirkung der wärme auf die organische welt betrachtet. Meteorologische Zeitschrift 1(21), 5–226 (1884)

6. Pusadee, T., Wongtamee, A., Rerkasem, B., Olsen, K.M., Jamjod, S.: Farmers drive genetic diversity of thai purple rice (oryza sativa l.) landraces. Economic Botany 73(1), 76–85 (2019)

7. Pathaichindachote, W., Panyawut, N., Sikaewtung, K., Patarapuwadol, S., Muangprom, A.: Genetic diversity and allelic frequency of selected thai and exotic rice germplasm using ssr markers. Rice Science 26(6), 393–403 (2019)

8. Kladmook, M., Kumchoo, T., Hongtrakul, V.: Genetic diversity analysis and subspecies classification of thailand rice landraces using dna markers. African Journal of Biotechnology 11(76), 14044–14053 (2012)

9. Reig-Valiente, J.L., Viruel, J., Sales, E., Marqués, L., Terol, J., Gut, M., Derdak, S., Talón, M., Domingo, C.: Genetic diversity and population structure of rice varieties cultivated in temperate regions. Rice 9(1), 58 (2016)

10. Pusadee, T., Schaal, B.A., Rerkasem, B., Jamjod, S.: Population structure of the primary gene pool of oryza sativa in thailand. Genetic Resources and Crop Evolution 60(1), 335–353 (2013)

11. Moonsap, P., Laksanavilat, N., Tasanasuwan, P., Kate-Ngam, S., Jantasuriyarat, C.: Assessment of genetic variation of 15 thai elite rice cultivars using indel markers. Crop Breeding and Applied Biotechnology 19(1), 15–21 (2019)

12. Zhao, K., Tung, C.-W., Eizenga, G.C., Wright, M.H., Ali, M.L., Price, A.H., Norton, G.J., Islam, M.R., Reynolds, A., Mezey, J., et al.: Genome-wide association mapping reveals a rich genetic architecture of complex traits in oryza sativa. Nature communications 2(1), 1–10 (2011)

13. Holsinger, K.E., Weir, B.S.: Genetics in geographically structured populations: defining, estimating and interpreting f st. Nature Reviews Genetics 10(9), 639 (2009)

14. Li, J.-Y., Wang, J., Zeigler, R.S.: The 3,000 rice genomes project: new opportunities and challenges for future rice research. Gigascience 3(1), 2047–217 (2014)

15. Mikami, K., Katagiri, T., Iuchi, S., Yamaguchi-Shinozaki, K., Shinozaki, K.: A gene encoding phosphatidylinositol-4-phosphate 5-kinase is induced by water stress and abscisic acid in arabidopsis thaliana. The Plant Journal 15(4), 563–568 (1998). doi:10.1046/j.1365-313X.1998.00227.x. https://onlinelibrary.wiley.com/doi/pdf/10.1046/j.1365-313X.1998.00227.x

16. Yamaguchi, T., Lee, D.Y., Miyao, A., Hirochika, H., An, G., Hirano, H.-Y.: Functional diversification of the two c-class mads box genes osmads3 and osmads58 in oryza sativa. The Plant Cell 18(1), 15–28 (2006)

17. Yun, D., Liang, W., Dreni, L., Yin, C., Zhou, Z., Kater, M.M., Zhang, D.: Osmads16 genetically interacts with osmads3 and osmads58 in specifying floral patterning in rice. Molecular Plant 6(3), 743–756 (2013). doi:10.1093/mp/sst003

18. Dreni, L., Pilatone, A., Yun, D., Erreni, S., Pajoro, A., Caporali, E., Zhang, D., Kater, M.M.: Functional Analysis of All AGAMOUS Subfamily Members in Rice Reveals Their Roles in Reproductive Organ Identity Determination and Meristem Determinacy. The Plant Cell 23(8), 2850–2863 (2011). doi:10.1105/tpc.111.087007. https://academic.oup.com/plcell/article-pdf/23/8/2850/36936516/plcell_v23_8_2850.pdf

19. Chen, R., Shen, L.-P., Wang, D.-H., Wang, F.-G., Zeng, H.-Y., Chen, Z.-S., Peng, Y.-B., Lin, Y.-N., Tang, X., Deng, M.-H., Yao, N., Luo, J.-C., Xu, Z.-H., Bai, S.-N.: A gene expression profiling of early rice stamen development that reveals inhibition of photosynthetic genes by osmads58. Molecular Plant 8(7), 1069–1089 (2015). doi:10.1016/j.molp.2015.02.004

20. Li, H., Liang, W., Hu, Y., Zhu, L., Yin, C., Xu, J., Dreni, L., Kater, M.M., Zhang, D.: Rice MADS6 Interacts with the Floral Homeotic Genes SUPERWOMAN1, MADS3, MADS58, MADS13, and DROOPING LEAF in Specifying Floral Organ Identities and Meristem Fate. The Plant Cell 23(7), 2536–2552 (2011). doi:10.1105/tpc.111.087262. https://academic.oup.com/plcell/article-pdf/23/7/2536/36108068/plcell_v23_7_2536.pdf

21. Zhang, H., Li, L., He, Y., Qin, Q., Chen, C., Wei, Z., Tan, X., Xie, K., Zhang, R., Hong, G., Li, J., Li, J., Yan, C., Yan, F., Li, Y., Chen, J., Sun, Z.: Distinct modes of manipulation of rice auxin response factor osarf17 by different plant rna viruses for infection. Proceedings of the National Academy of Sciences 117(16), 9112–9121 (2020). doi:10.1073/pnas.1918254117. https://www.pnas.org/content/117/16/9112.full.pdf

22. Chen, S.-H., Zhou, L.-J., Xu, P., Xue, H.-W.: Spoc domain-containing protein leaf inclination3 interacts with lip1 to regulate rice leaf inclination through auxin signaling. PLOS Genetics 14(11), 1–19 (2018). doi:10.1371/journal.pgen.1007829

23. Li, Y., Li, J., Chen, Z., Wei, Y., Qi, Y., Wu, C.: Osmir167a-targeted auxin response factors modulate tiller angle via fine-tuning auxin distribution in rice. Plant Biotechnology Journal 18(10), 2015–2026 (2020). doi:10.1111/pbi.13360. https://onlinelibrary.wiley.com/doi/pdf/10.1111/pbi.13360

24. Wu, L., Cui, Y., Xu, Z., Xu, Q.: Identification of multiple grain shape-related loci in rice using bulked segregant analysis with high-throughput sequencing. Frontiers in Plant Science 11, 303 (2020). doi:10.3389/fpls.2020.00303

25. PRJNA753279-Thai Rice Genotyping Project. https://dataview.ncbi.nlm.nih.gov/object/PRJNA753279?reviewer=i3jrmvv07t4g6n268gsu3ub5q4. Accessed: 2021-09-15

26. Li, H., Handsaker, B., Wysoker, A., Fennell, T., Ruan, J., Homer, N., Marth, G., Abecasis, G., Durbin, R.: The sequence alignment/map format and samtools. Bioinformatics 25(16), 2078–2079 (2009)

27. Li, H.: Aligning sequence reads, clone sequences and assembly contigs with bwa-mem. arXiv preprint 1303.3997 (2013)

28. McKenna, A., Hanna, M., Banks, E., Sivachenko, A., Cibulskis, K., Kernytsky, A., Garimella, K., Altshuler, D., Gabriel, S., Daly, M., et al.: The genome analysis toolkit: a mapreduce framework for analyzing next-generation dna sequencing data. Genome research 20(9), 1297–1303 (2010)

29. Bradbury, P.J., Zhang, Z., Kroon, D.E., Casstevens, T.M., Ramdoss, Y., Buckler, E.S.: TASSEL: software for association mapping of complex traits in diverse samples. Bioinformatics 23(19), 2633–2635 (2007). doi:10.1093/bioinformatics/btm308. http://oup.prod.sis.lan/bioinformatics/article-pdf/23/19/2633/451862/btm308.pdf

30. Alexander, D.H., Novembre, J., Lange, K.: Fast model-based estimation of ancestry in unrelated individuals. Genome research 19(9), 1655–1664 (2009)

31. Forgy, E.W.: Cluster analysis of multivariate data: efficiency versus interpretability of classifications. Biometrics 21, 768–769 (1965)

32. R Development Core Team, R., et al.: R: A language and environment for statistical computing. R foundation for statistical computing Vienna, Austria (2011)

33. Efron, B.: Bootstrap Methods: Another Look at the Jackknife, pp. 569–593. Springer, New York, NY (1992). doi:10.1007/978-1-4612-4380-941

34. Saitou, N., Nei, M.: The neighbor-joining method: a new method for reconstructing phylogenetic trees. Molecular Biology and Evolution 4(4), 406–425 (1987). doi:10.1093/oxfordjournals.molbev.a040454. http://oup.prod.sis.lan/mbe/article-pdf/4/4/406/11167444/7sait.pdf

35. Amornbunchornvej, C., Surasvadi, N., Plangprasopchok, A., Thajchayapong, S.: A nonparametric framework for inferring orders of categorical data from category-real pairs. Heliyon 6(11), 05435 (2020). doi:10.1016/j.heliyon.2020.e05435

36. Breiman, L.: Random forests. Machine learning 45(1), 5–32 (2001)

37. Liaw, A., Wiener, M.: Classification and regression by randomforest. R News 2(3), 18–22 (2002)

38. Kuhn, M.: Caret: Classification and Regression Training. (2020). R package version 6.0-86. https://CRAN.R-project.org/package=caret

39. Allen, D.M.: The relationship between variable selection and data agumentation and a method for prediction. Technometrics 16(1), 125–127 (1974). doi:10.1080/00401706.1974.10489157. https://www.tandfonline.com/doi/pdf/10.1080/00401706.1974.10489157

